## Supplementary_File_2 for "Hierarchical architecture of dopaminergic circuits enables second-order conditioning in *Drosophila*"

**KEY RESOURCES TABLE**

| **REAGENT or RESOURCE** | **SOURCE** | **IDENTIFIER** |
| --- | --- | --- |
| **Antibodies** | | |
| rabbit polyclonal anti-GFP (1:1000) | Invitrogen | A11122  RRID:AB_221569 |
| mouse monoclonal anti-Brp (1:30) | *Developmental Studies Hybridoma Bank* | nc82  RRID:AB_2341866 |
| mouse monoclonal anti-ChAT (1:50) | *Developmental Studies Hybridoma Bank* | ChAT4B1  RRID:AB_528122 |
| rabbit monoclonal anti-HA-Tag (1:300) | Cell Signaling Technology | C29F4; #3724  RRID:AB_10693385 |
| rat anti-FLAG (1:200) | Novus Biologicals | NBP1-06712  RRID:AB_1625981 |
| mouse monoclonal anti-V5-TAG Dylight-549 (1:500) | Bio-Rad | MCA2894D549GA  RRID:AB_10845946 |
| goat polyclonal anti-mous IgG(H&L) AlexaFluor-568 (1:400) | Invitrogen | A11031  RRID:AB_144696 |
| goat polyclonal anti-rabbit IgG(H&L) AlexaFluor-488 (1:800) | Invitrogen | A11034  RRID:AB_2576217 |
| donkey polyclonal anti-mouse IgG(H&L) AlexaFluor-488 conjugated (1:400) | Jackson Immuno Research Labs | 715-545-151  RRID:AB_2341099 |
| donkey polyclonal anti-rabbit IgG(H&L) AlexaFluor-594 (1:500) | Jackson Immuno Research Labs | 711-585-152  RRID:AB_2340621 |
| donkey polyclonal anti-rat IgG(H&L) AlexaFluor-647 (1:300) | Jackson Immuno Research Labs | 712-605-153  RRID:AB_2340694 |
| goat polyclonal anti-Mouse IgG (H&L) ATTO 647N (1:100) | ROCKLAND | 610-156-121  RRID:AB_10894200 |
| goat polyclonal anti-rabbit IgG (H+L) Alexa Fluor 568  (1:1000) | Invitrogen | A-11036  RRID:AB_10563566 |
| **Chemicals** | | |
| 3-Octanol | Sigma-Aldrich | 218405 |
| 4-Methylcyclohexanol | VWR | AAA16734-AD |
| Pentyl acetate | Sigma-Aldrich | 109584 |
| Ethyl lactate | Sigma-Aldrich | W244015 |
| Paraffin oil | Sigma-Aldrich | 18512 |
| **Deposited data** | | |
| Confocal microscopy images of split-GAL4 drivers | This paper | <http://flweb.janelia.org/cgi-bin/splitgal4.cgi> |
| **Experimental models: Organisms/strains** | | |
| Canton S | Martin Heisenberg | N.A. |
| *20xUAS-CsChrimson-mVenus attP18* | Klapoetke et al., 2014; PMID: 24509633 | N.A. |
| *10XUAS-Chrimson88-tdTomato attP1* | Klapoetke et al., 2014; PMID: 24509633 | N.A. |
| 13XLexAop2-IVS-ChrimsonR-mVenus-p10 attP18 | Vivek Jayaraman | N.A. |
| 20XUAS-syn21-mScarlet-opt-p10 su(Hw)attp8 | Glenn Turner | N.A. |
| *pJFRC200-10xUAS-IVS-myr::smGFP-HA in attP18* | Nern et al.,2015; PMID: 25964354 | N.A. |
| *pJFRC225-5xUAS-IVS-myr::smGFP-FLAG in VK00005* | Nern et al.,2015; PMID: 25964354 | N.A. |
| *pBPhsFlp2::PEST in attP3* | Nern et al.,2015; PMID: 25964354 | N.A. |
| *pJFRC201-10XUAS-FRT>STOP>FRT-myr::smGFP-HA in VK0005* | Nern et al.,2015; PMID: 25964354 | N.A. |
| *pJFRC240-10XUAS-FRT>STOP>FRT-myr::smGFP-V5-THS-10XUAS-FRT>STOP>FRT-myr::smGFP-FLAG_in_su(Hw)attP1* | Nern et al.,2015; PMID: 25964354 | N.A. |
| *LexAop2-DA2m VK00005* | Sun et al., 2020; PMID: 33087905 | N.A. |
| MB043-split-LexA | This paper | N.A. |
| *empty-split-GAL4 (p65ADZp attP40, ZpGAL4DBD attP2)* | Seeds et al., 2014; PMID: 25139955 | N.A. |
| MB032B split-GAL4 | Aso et al., 2014a; PMID: 25535793 | N.A. |
| MB043C split-GAL4 | Aso et al., 2014a; PMID: 25535793 | N.A. |
| MB109B split-GAL4 | Aso et al., 2014a; PMID: 25535793 | N.A. |
| MB213B split-GAL4 | Aso et al., 2014a; PMID: 25535793 | N.A. |
| MB315C split-GAL4 | Aso et al., 2014a; PMID: 25535793 | N.A. |
| SS33917 split-GAL4 | This paper | N.A. |
| SS45234 split-GAL4 | This paper | N.A. |
| SS67221 split-GAL4 | This paper | N.A. |
| UAS-TeNT | Keller et al., 2002: PMID: 11810637 | N.A. |
| **Software and algorithms** | | |
| ImageJ and Fiji | NIH  Schneider et al., 2012 | <https://imagej.nih.gov/ij/>  <http://fiji.sc/> |
| MATLAB | MathWorks | <https://www.mathworks.com/> |
| Adobe Illustrator CC | Adobe Systems | <https://www.adobe.com/products/illustrator.html> |
| GraphPad Prism 9 | GraphPad Software | <https://www.graphpad.com/scientific-software/prism/> |
| Python | Python Software Foundation | <https://www.python.org/> |
| neuPrint | HHMI Janelia | <https://doi.org/10.25378/janelia.12818645.v1> |
| Cytoscape | (Shannon et al., 2003) | https://cytoscape.org/ |
| NeuTu | [Zhao et al., 2018](https://elifesciences.org/articles/62576#bib209) | https://github.com/janelia-flyem/NeuTu |
| ScanImage | Vidrio Technologies | https://vidriotechnologies.com/ |
| VVDveiwer | HHMI Janelia | <https://github.com/takashi310/VVD_Viewer> |
| **Other** | | |
| Grade 3MM Chr Blotting Paper | Whatmann | 3030-335 |
| mass flow controller | Alicat | MCW-200SCCM-D |
| optogenetic olfactory arena | Aso and Rubin 2016 | N.A. |
